# Structure-function analysis of distant diphtheria toxin homologs reveals insights into host adaptation

**DOI:** 10.1101/2021.08.10.455837

**Authors:** Seiji N. Sugiman-Marangos, Shivneet K. Gill, Michael J. Mansfield, Kathleen E. Orrell, Andrew C. Doxey, Roman A. Melnyk

**Affiliations:** Molecular Medicine Program, The Hospital for Sick Children Research Institute, 686 Bay Street Toronto, ON, Canada, M5G 0A4; The Hospital for Sick Children Research Institute, 686 Bay Street Toronto, ON, Canada, M5G 0A4; Department of Biochemistry, University of Toronto, Toronto, ON, Canada, M5S1A8; Genomics and Regulatory Systems Unit, Okinawa Institute of Science and Technology Graduate University, Onna, Okinawa, Japan; Deparment of Biology, University of Waterloo, ON, Canada

## Abstract

Diphtheria toxin (DT) is the archetype of bacterial exotoxins implicated in human diseases and has played a central role in defining the field of toxinology since its discovery in 1888. Despite being one of the most extensively characterized bacterial toxins, the origins and molecular evolution of DT host specialization remain unknown. Here, we determined high-resolution structures of two recently discovered distant homologs of DT. These DT-like proteins from non-human associated *Streptomyces albireticuli* (17% identity to DT) and *Seinonella peptonophila* (20% identity to DT) display remarkable structural similarity to DT enabling a comparative investigation into DT’s unique toxicity toward mammalian cells. We find that the individual domains of DT-like toxins retain two critical features of DT’s activity: full catalytic function and ability to translocate across mammalian cell membranes. However, we show that receptor-binding, pH-dependent pore-formation and proteolytic release of the cytotoxic enzyme into the cytosol are not optimized for human cell physiology and thus unable to efficiently deliver the cytotoxic cargo into human hosts. Our work provides structural insights into DT’s evolutionary history, and implies key transitions required for the emergence of human-specificity of a major bacterial exotoxin with an important history in human disease.

## Introduction

The disease symptoms that accompany many bacterial infections are caused by the actions of pathogen-derived proteins known as exotoxins. Since the discovery of the first bacterial exotoxin diphtheria toxin (DT) in 1888, as the cause of major symptoms of diphtheria, the field of toxinology has made considerable progress in understanding the structures and functions of these remarkably toxic proteins. The most potent toxins, such as botulinum toxin, tetanus toxin and diphtheria toxin, despite having no discernable sequence similarities, all shared a common architecture of three functional units: a cytotoxic enzymatic domain (C), a translocation domain (T), and a receptor binding domain (R). Nearly a century of research on the structure, function, and mechanisms of action of DT has uncovered insights into how this multifunctional protein binds, penetrates and intoxicates host cells to ultimately cause disease (Figure 1A) [1]. Our knowledge of the evolutionary origins and molecular ancestry of DT, by contrast, remains poor due in large part to the lack of characterized relatives of DT. Traditional identification of novel toxins involves clinical isolates from bacterial infections of humans or livestock. As a result, the field of toxinology has been human-centric, complicating investigations of toxin evolution since ancestrally related proteins need not be related to pathogenesis in human hosts. Advances in rapid genome sequencing technologies have provided access to vast quantities of data which has led to the discovery of a variety of interesting bacterial toxins that might not otherwise have been studied[2]. This includes the identification of new members of the botulinum neurotoxin superfamily[3], anthrax toxin family[4], large clostridial toxin family[5] and our recent identification of DT homologs outside of *Corynebacterium* [6].

**Figure 1:**
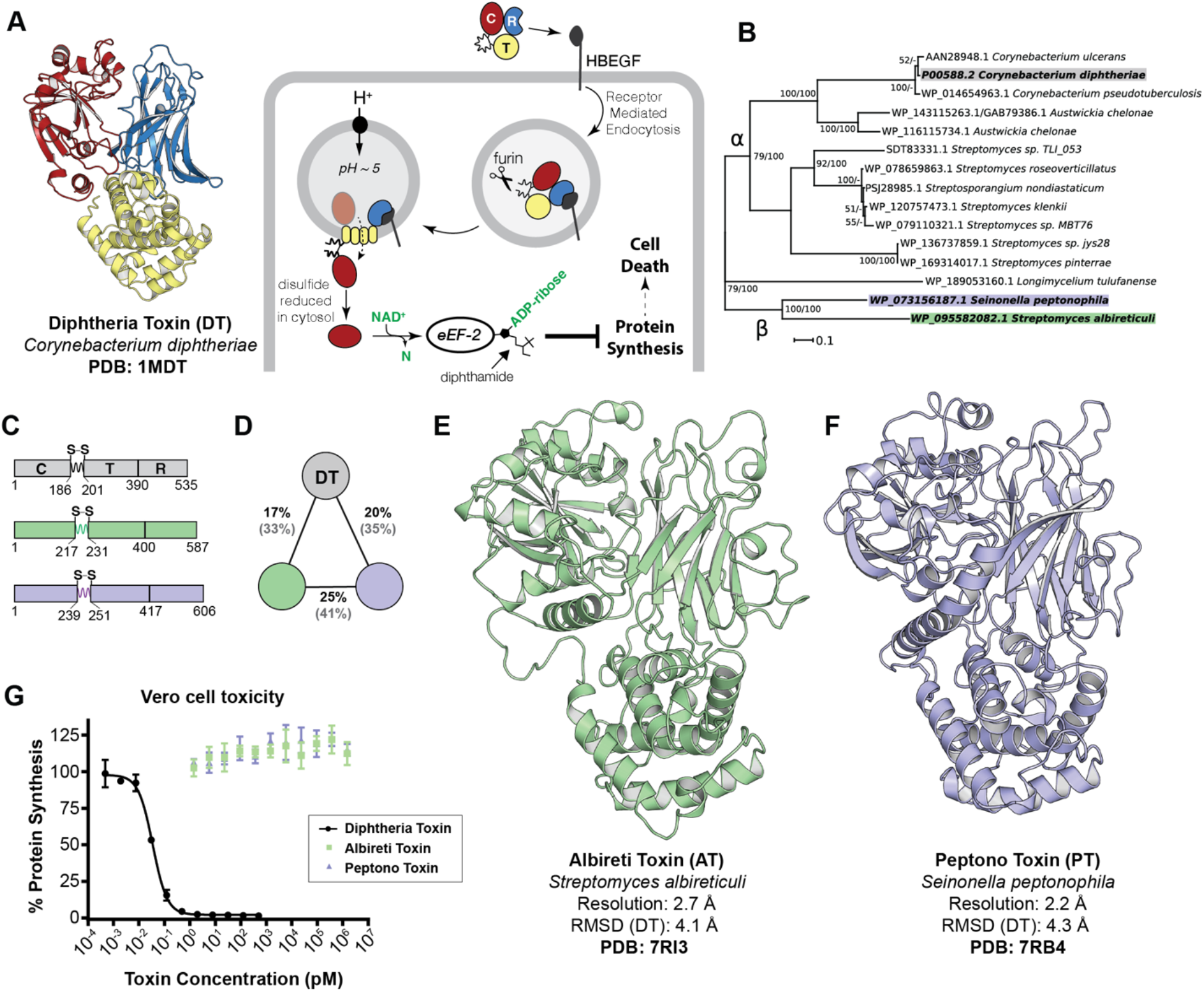
Structures and functions of remote diphtheria toxin homologs. (A) DT intoxication pathway. Briefly, the C-terminal <u>**R**</u>eceptor binding domain of DT (blue) triggers receptor mediated endocytosis upon binding its receptor HBEGF. In the endosome, a furin-like protease cleaves between the <u>**C**</u>ytotoxic domain (red) and the <u>**T**</u>ranslocation domain (yellow) which remain tethered by an intramolecular disulfide bond. Acidification due to endosomal maturation results in conformational changes in the T domain which then inserts into the membrane, facilitating the escape of the C domain into the cytosol. The reducing environment of the cytosol reduces the disulfide bond, freeing the C domain to ADP-ribosylate its target eEF-2. ADP-ribosylation of the diphthamide residue on eEF-2 inactivates it, leading to protein synthesis inhibition and cell death. (B) Phylogenetic tree of DT-like proteins sub-divided into α and β-clades. DT and DT-like proteins from *S. albireticuli* (AT) and *S. peptonophila* (PT) are highlighted in grey, green and blue, respectively. Clade support values correspond to maximum likelihood bootstrap values from RAxML and posterior probability percentages from MrBayes. (C) Domain architecture and residue boundaries of DT, AT, and PT. (D) Three-way sequence identity and similarity (brackets) between DT, AT, and PT. (E) Crystal structure of AT solved by SAD-phasing and refined to R and R_free_ values of 20.2% and 24.8%. The four monomers in the ASU align to each other with RMSDs of <1 .0 Å. A complete model was built in each of the four chains, except for 4-5 residues from the N-terminus and 0-10 residues from the loop joining the catalytic and translocation domains. Shown is the most complete of the four chains, C. (F) Crystal structure of Peptono Toxin solved by SAD-phasing and refined to R and R_free_ values of 20.2% and 22.5%. A complete model was built for the monomer in the ASU, apart from the first three N-terminal residues. (G) Dose titration of DT, AT, and PT on DT-sensitive Vero cells (mean ± SD, n=3).

The discovery of DT homologs has provided a unique opportunity to study the origins of DT and potentially understand the determinants of its toxicity toward humans. Until this discovery, the only known homologs (from *C. ulcerans* and *C. pseudotuberculosis*) were too closely related (shared ≥94% sequence identity) to provide meaningful insights into the factors contributing to DT’s host specificity. This growing lineage of DT-like proteins from non-human associated organisms (Figure 1B) contain all the necessary domains to constitute legitimate exotoxins (Figure 1C). Studying these distant homologs, which are found in organisms that have not been implicated as human pathogens, provides an opportunity to better understand the determinants of DT’s toxicity in humans. We therefore characterized the DT-like homologs from *Streptomyces albireticuli* and *Seinonella peptonophila*, representing the most divergent homologs from DT (17% and 20% sequence identity to DT, respectively). Of note, while they group together in this growing family of DT-like proteins, they are nearly as dissimilar to one another (25% sequence identity) as they are to DT (Figure 1D). In this study, we report the first crystal structures of DT-like proteins and carry out a rigorous characterization of each functional domain. We identify several features shared amongst the three proteins and outline key structural and functional differences relevant to DT’s host specificity.

## Results

### Structural elucidation of DT-like proteins from *S. albireticuli and S. peptonophila*

To structurally characterize the DT-like proteins from *S. albireticuli* and *S peptonophila*, their gene sequences (WP_095582082.1 and WP_073156187.1 respectively) were first synthesized, incorporating codons that were optimized for expression in *E. coli*. Crystals of affinity purified proteins were grown by hanging-drop vapor diffusion and diffraction data was collected on the AMX beamline at NSLSII. Neither homolog could be solved by molecular replacement with the structure of DT, and therefore both required experimental phases derived from single-wavelength anomalous dispersion (SAD) datasets to be solved. Experimental phase for the DT-like protein from *S. albireticuli* was obtained from a 2.7 Å SAD dataset using selenomethionine (SeMet) derivatized protein crystals. As crystals of the DT homolog from *S. peptonophila* were grown in a condition containing potassium iodide, a 2.2 Å dataset was collected with native protein crystals at a wavelength of 2.0 Å to maximize the anomalous signal from iodide. The structure was subsequently solved using iodide-SAD phasing. A complete list of diffraction data and model refinement statistics for both structures can be found in Supplemental Table 1.

### Remote DT-like proteins display structural homology to DT

Both DT-like proteins, despite their low identity to DT and to each other (Figure 1D), were found to adopt the characteristic three domain Y-shaped architecture of DT (Figure 1E-F). Moreover, each of the individual domains preserve their respective folds: the central split β-sheet motif in the catalytic domain characteristic of the ADP-ribosyl transferase (ART) superfamily; the three-layered α-helical translocation domain; and, the Ig-like jelly-roll fold in the receptor-binding domain. With respect to insertions and deletions, the T-domain appears to be the most structurally conserved of the three domains, while both the C and R-domains contain several additional secondary-structure features not present in DT. Structural alignments performed with the DALI server [7] calculate the overall RMSD of the structures to DT as 4.1 Å (*S. albireticuli* - 468/583 residues) and 4.3 Å (*S. peptonophila* - 457/603 residues). Additionally, both DT homologs maintain the pinched disulfide loop at the junction between the C- and T-domains. Given these structural similarities, and for purposes of clarity and simplicity, we refer to these two DT homologs hereafter as Albireti Toxin (AT) and Peptono Toxin (PT).

### AT and PT holotoxins are non-toxic to DT-sensitive Vero Cells

The striking structural conservation despite low sequence conservation of AT and PT with DT (Figure 1A,E,F) prompted us to explore their potential to function as toxins. AT and PT were added to Vero cells and incubated overnight; protein synthesis was measured using a nanoluciferase (nLucP) based assay[8] following an overnight (18 hour) incubation. Vero cells express high levels of the DT-receptor[9], HBEGF, and are therefore particularly sensitive to DT. Under these conditions DT enters cells and inhibits protein synthesis with an EC_50_ of ∼0.15 pM (Figure 1G). By striking contrast, we found that AT and PT do not display any measurable toxicity on Vero cells even at µM concentrations (Figure 1G). The observation that these DT homologs exhibited no cellular toxicity on Vero cells while having remarkable structural homology to DT, provided an unprecedented opportunity to elucidate the determinants of toxicity for DT. To this end, we carried out an in-depth, domain-by-domain, functional/structural comparison with the goal of better understanding DT’s potency in human hosts.

### AT and PT catalytic domains are potent modifiers of eEF-2

Structurally, the catalytic domains from AT and PT contain many of the hallmarks of a functional ADP-ribosyltransferase. Both AT_C_ and PT_C_ contain the characteristic split β-sheet (β1-5-6-7-3 and β2-4-8) at the core of the ART fold (Figure 2A) shared amongst other ADP-ribosylating toxins (Supplemental Figure 1). Moreover, within the putative active sites, two key active site residues, Y54 and E148, are absolutely conserved and structurally equivalent in the two DT homologs (Figure 2B). To evaluate whether the individual catalytic moieties of AT and PT had cytotoxic activity if introduced into cells, we generated domain-swapped chimeras between DT and AT_C_ or PT_C_ and determined their toxicity on DT-sensitive cells (Figure 2C). In each DT chimera, DT_C_ was removed and replaced with the corresponding domain from the homolog, e.g., in DT(AT_C_), AT_1-216_ fused to DT_186-535_. A full description of domain boundaries used in DT chimeras is outlined in Supplemental Figure 2. Despite AT and PT holotoxins not displaying any toxicity on cells up to μM concentrations (Figure 1G), DT(AT_C_) and DT(PT_C_) inhibited protein synthesis with EC_50_’s of 1.5 and 22 pM, respectively (Figure 2C), indicating that AT_C_ and PT_C_ are in fact highly toxic if delivered into cells via DT_T_ and DT_R_.

**Figure 2.**
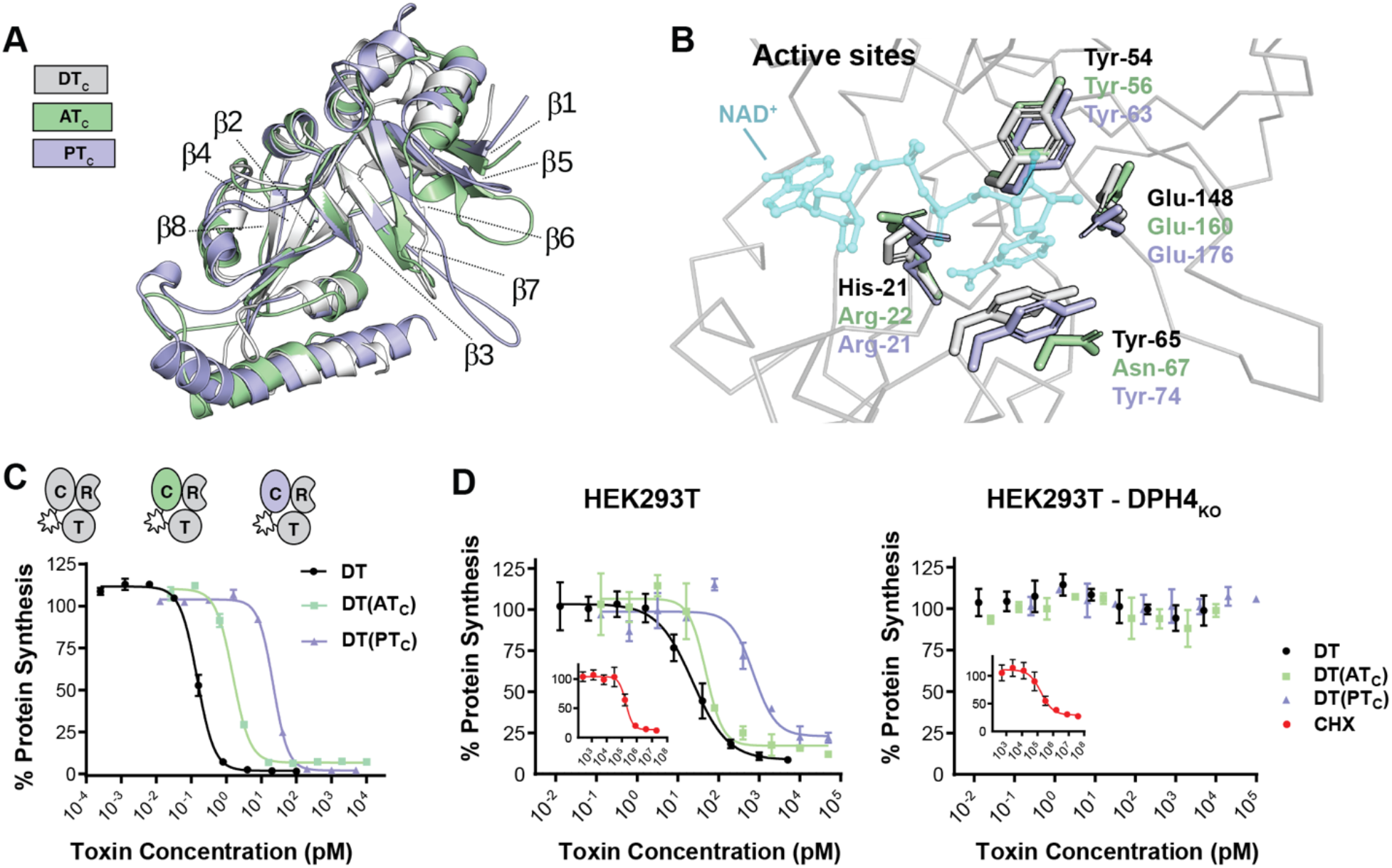
The catalytic moieties from AT and PT are cytotoxic. (A) Overlay of C-domains highlighting the conserved split β-sheet characteristic of ART fold proteins. (B) Active site residue conservation. DT residues important for NAD^+^ binding and catalysis and corresponding residues in AT and PT are represented in stick form. NAD^+^ represented in partial transparency is modeled from PDB entry 1TOX for reference. (C) Dose titration of DT and DT C-domain chimeras DT(AT_C_), DT(PT_C_) on Vero cells (mean ± SD, n=3). (D) Dose titration of DT, DT(AT_C_), and DT(PT_C_) on HEK293T and HEK293T DPH4_KO_ cells (mean ± SD, n=3). Titration of protein synthesis inhibitor cycloheximide (CHX) is shown in red (inset).

Next, to determine whether the homologs induce cytotoxicity by targeting the same eEF-2 residue as DT (diphthamide), we examined a CRISPR knockout cell-line unable to synthesize diphthamide (HEK239T DPH4_KO_). Wild-type and DPH4 _KO_ cells were treated with increasing concentrations of DT, DT(AT_C_), DT(PT_C_) or a chemical inhibitor of protein synthesis (cycloheximide), and protein synthesis was assessed following overnight incubation (Figure 2D). While both cell types were sensitive to cycloheximide, only the wild-type HEK293T cells were sensitive to DT and the catalytic domain chimeras. DPH4 _KO_ cells were unaffected by any of the toxins. The observed strict dependence on diphthamide biosynthesis demonstrates that AT_C_ and PT_C_ induce profound cytotoxicity through a DT_C_-like mechanism if introduced into cells.

### AT and PT lack the canonical protease activation site of DT

Of the seven homologs we identified previously [6], AT and PT were the only two lacking the canonical ‘RXXR’ furin cleavage motif in the loop joining the putative catalytic fragments with the downstream putative translocation and receptor binding fragments (Figure 3A). Of note, however, AT and PT contain both Cys residues (C217/C231 and C239/C251, respectively) that bound this region, and the crystal structures confirm formation of the inter-fragment disulfide bonds (Figure 3A). To determine whether the intervening loop could still be recognized and cleaved by another cellular protease, DT, AT, and PT were incubated with total cell lysate from Vero cells. To evaluate whether cleavage occurred, toxins were separated by SDS-PAGE under reducing conditions. Whereas DT was processed into its two fragments following overnight incubation with lysate, neither of the homologs showed any significant processing (Figure 3B). This was unexpected given that it is generally accepted that cleavage of the disulfide loop between the two fragments is a crucial step in the DT intoxication pathway [10, 11]. To determine the functional effect of the reduced activation by cleavage, we replaced the residues subtended by C186 and C201 from DT with the corresponding residues from AT and PT and assessed their toxicity. The DT(AT_F_) and DT(PT_F_) chimeras were toxic to Vero cells each with EC_50_’s of 22 pM (Figure 3C), or approximately two orders-of-magnitude less potent than DT. Although DT(AT_F_) and DT(PT_F_) are less toxic than DT, if release of the catalytic domain into the cytosol is absolutely required for toxicity, these constructs would be expected to be completely non-toxic.

**Figure 3.**
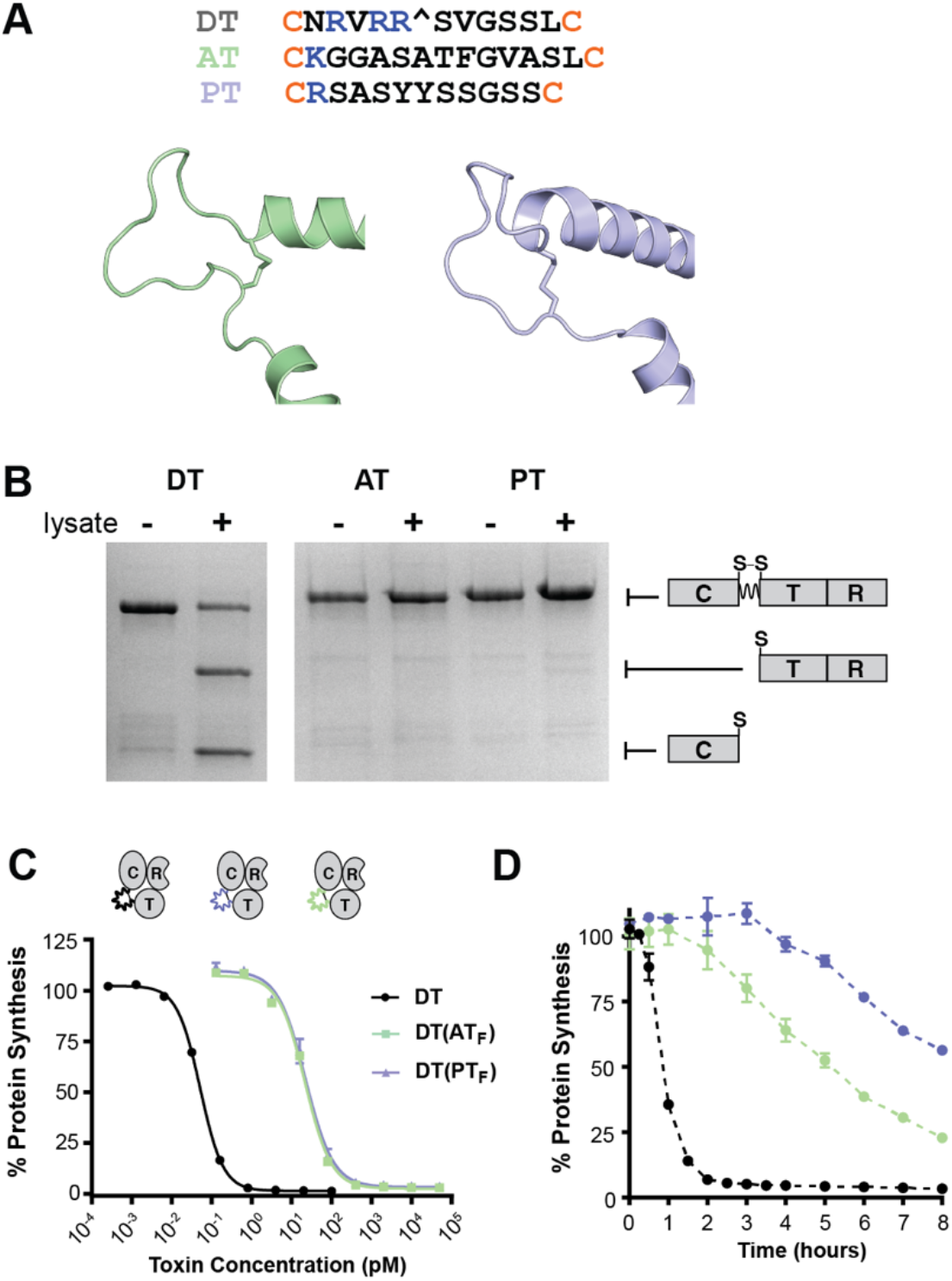
AT and PT are not efficiently processed. (A) Primary amino acid sequences of loops linking C and T domains in DT, AT, and PT. Loop structures of AT (green) and PT (blue) demonstrating intramolecular disulfide bond formation. (B) SDS-PAGE analysis of holotoxin processing by Vero cell lysate. (C) Dose titration of DT and DT activation loop chimeras DT(AT_F_), DT(PT_F_) on Vero cells (mean ± SD, n=3). (D) Time-course of protein synthesis inhibition in Vero cells treated with 10 nM of DT, DT(AT_F_), and DT(PT_F_) (mean ± SD, n=3).

To further investigate the functional consequences of having an ostensibly non-cleavable site in place of the furin processing site in DT, Vero cells were incubated with 10 nM of DT, DT(AT_F_) or DT(PT_F_), a concentration which after overnight incubation results in complete cell death, and protein synthesis was evaluated at regular intervals for a total of 8 hours. Whereas cells treated with DT experienced a rapid onset of protein synthesis inhibition 1 hour after toxin administration, cells treated with DT(AT_F_) and DT(PT_F_) demonstrated slower kinetics of inhibition of protein synthesis (Figure 3D), highlighting a potential role for the proteolytic processing event in accelerating the intoxication process.

### AT and PT translocation domains require lower pH for membrane insertion

The T domain is arguably the most specialized domain of DT: the acidic environment of the early endosome induces partial unfolding and formation of a molten globule state followed by insertion of at least two marginally hydrophobic segments into the endosomal membrane to form a pore that catalyzes delivery of the toxic C domain into the cytosol. Structurally, both AT_T_ and PT_T_ maintain the same three-layered helical architecture as DT_T_, only lacking structural equivalents to TH2 (PT) and TH4 (AT and PT) (Figure 4A). Notably, the hydrophobic helices that make up the “double-dagger” motif (TH5-9)[12] are particularly well conserved structurally, consistent with the disproportionate sequence conservation of the first half (TH1-4) vs the second half (TH5-9) of the T domains (AT - 13% cf 21%, PT - 25% cf 39%). The loop joining TH8 and TH9 is thought to initiate insertion into the endosomal membrane upon partial protonation of E349 and D352 during endosomal acidification[12-14] (Figure 4B). In addition to the two negatively charged residues, P345 is thought to play an important role in mediating membrane insertion of the T domain [15, 16]. Both AT and PT contain an equivalent Pro residue at the C-terminus of TH8 that breaks the helix (Figure 4B). The loop in AT contains only one negatively charged residue which sits within the first turn of TH9 rather than in the loop itself, while PT contains equivalents to both DT_E349_ and DT_D352_ (Figure 4B).

**Figure 4.**
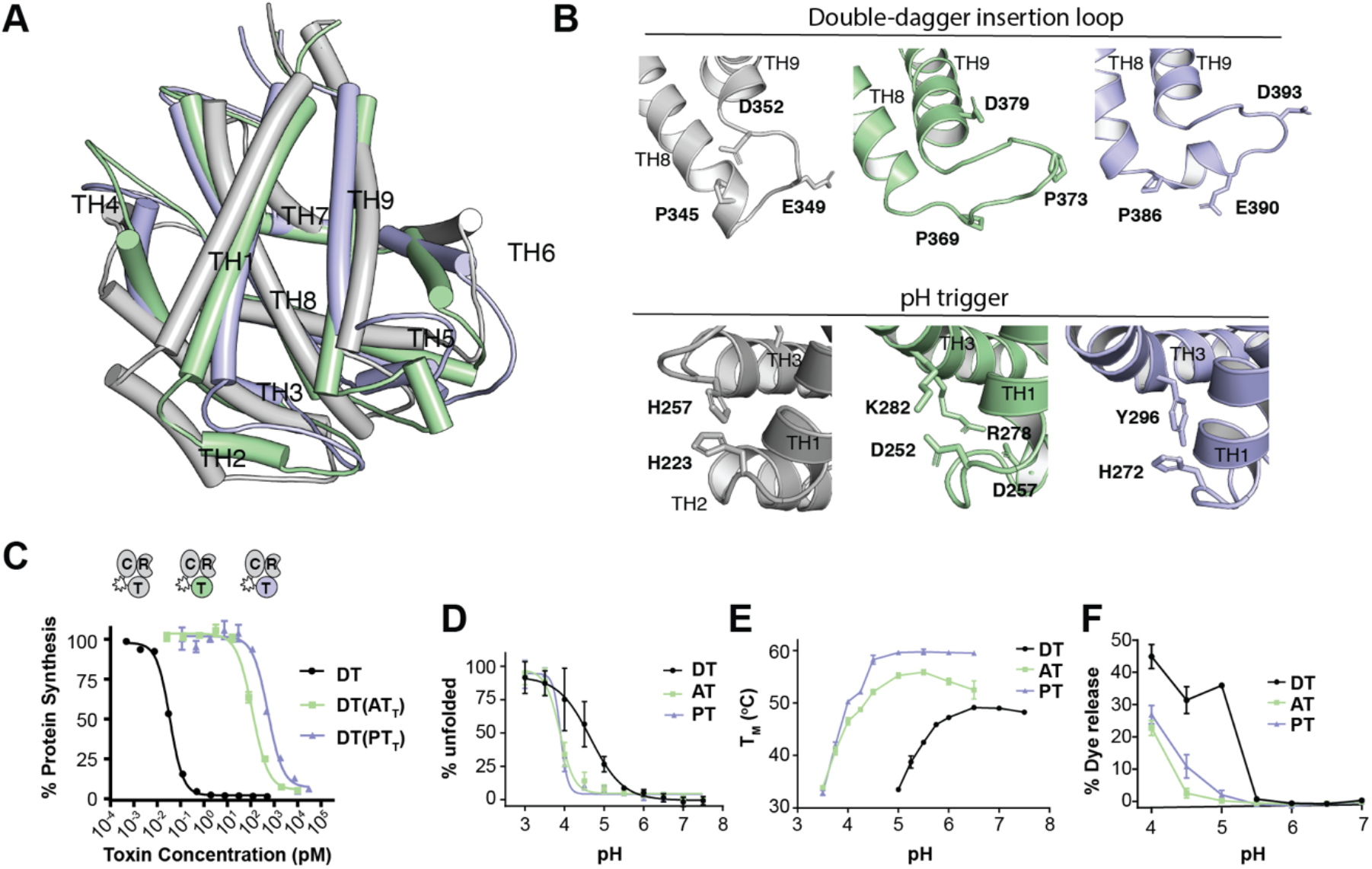
AT and PT have functional translocases with altered pH dependence. (A) Overlay of T-domains structures (AT: 20% sequence identity, RMSD 3.0 Å; PT: 31% sequence identity, RMSD 3.1 Å). (B) Residues of interest in the TH8/TH9 loop involved in initial membrane insertion of DT_T_ (top). AT and PT contain an equivalent to DT_P345_ (AT_P369_ and PT_P386_). The loop in AT contains a three-residue insertion and an additional Pro residue within the loop (AT_P373_), and only one negatively charged residue (AT_D379_). PT_P386_ causes a 90° kink in TH8 which continues for a full turn before terminating with a Glu residue (PT_E390_). Residues of interest at the junction of TH1, TH2, and TH3 believed to play an important role in early destabilization of DT_T_ (bottom). AT_T_ contains two salt-bridges (D252/R282, D257/K278) in lieu of His residues in this junction (C) Toxicity of chimeric toxins on Vero cells. Dose titration of DT and DT T-domain chimeras DT(AT_T_), and DT(PT_T_) on Vero cells (mean ± SD, n=3). (D) pH dependent TNS analysis. While DT begins to unfold at pH <6 (pKa ∼4.7), AT and PT begin unfolding at pH <5 (pKa ∼3.9). (E) pH dependent thermal denaturation by DSF. The transition temperature (T_m_) of DT begins to decrease below pH 6 while AT and PT do not begin to decrease until pH values below pH 5. (F) pH dependent dye leakage from liposomes loaded with fluorophore quencher pair (HPTS/DPX). DT exhibits a spike of activity upon exposure to pH 5, while AT and PT are only active at pH 4, a full pH unit lower than DT.

To test whether the T-domains from AT and PT could function as protein translocases, as above we generated chimeric constructs with DT in which the T domain of DT was replaced with the T domain from AT or PT. DT(AT_T_) and DT(PT_T_) displayed toxicity to Vero cells with EC_50_’s of 117 and 535 pM, respectively (Figure 4C), demonstrating that both AT_T_ and PT_T_ are functional translocases, albeit with significantly decreased apparent efficiency as compared with DT_T_. A potential clue to explain the reduced ability of AT_T_ and PT_T_ to act as translocases in Vero cells came from a closer look at the key residues implicated in the pH-trigger for DT (reviewed in [17]). Of the six His residues in DT_T_, two have been identified as playing particularly crucial roles in initial unfolding events: H223 and H257, located in the loops between TH1/2 and TH3/4, respectively (Figure 4B). Protonation of H257 is thought to destabilize the first four helices of DT_T_[18], exposing the hydrophobic helices (TH5-8) and thereby promoting their interaction with the membrane [19]. H223 has been proposed to act as a “safety latch” to prevent premature unfolding [20]. Remarkably, the T-domain of AT lacks His residues in the T domain entirely. Instead, this region (TH1-4) is stabilized by two salt-bridges between residues in TH1/TH3 and TH2/TH3 (Figure 4B). The T domain of PT contains three His residues: H272, H276, and H364. Two of these residues are structurally equivalent with His residues in DT: PT_H272_↔DT_H223_ and PT_H364_↔DT_H322_. Rather than forming π-stacking interactions with a second His residue as in DT, PT_H272_ forms an edge-to-face π – π interaction with PT_Y296_ (Figure 4B).

Based on differences in the pH-sensitive determinants for DT, we posited that AT and PT may not be optimized to form functional translocases at early endosome pHs. To investigate this, we measured the pH-dependent unfolding and pore-formation of AT and PT using a TNS-fluorescence unfolding assay, differential scanning fluorescence and a dye-release assay of pore-formation. Consistent with observed differences in residues implicated in the pH-sensor in DT, we found in all three assays that AT and PT required a lower pH to unfold and form membrane-inserted pores (Figure 4D-F), suggesting that these translocases are not optimized to form functional translocases in early endosomes.

### AT_R_ and PT_R_ share the same structural fold as DT_R_ but do not bind hHBEGF

Of the three domains making up the holotoxin, the receptor-binding domains of AT and PT have the lowest sequence conservation with DT (16% and 13% sequence identity, respectively). We previously noted that these regions are highly divergent in sequence and were not recognizable using existing DT_R_ domain models [6]. Nevertheless, despite marginal sequence identity with DT_R_, the X-ray crystal structures reveal that AT_R_ and PT_R_ are in fact highly similar to DT_R_, maintaining the core β-sandwich motif (β1-2-3-8-5 and β6-7-9-10) present in DT_R_ (Figure 5A). Overall, AT_R_ and PT_R_ appear to be less compact than DT_R_, each containing ∼30 additional residues distributed amongst the loop regions joining secondary structure elements in the β-sandwich. Most of these residues are contained within two insertions present in both AT and PT: a 10 amino acid Ω-loop (7 amino acids in PT) and a 16 amino acid insertion in the loop joining β9 and β10. In DT_R_, β9 and β10 contribute a significant proportion of the HBEGF binding surface. In the bound state, the distal end of the DT_R_ β9-10 hairpin bends inward forming a concave surface like a grasping hand, breaking the continuity of the strands (Figure 5A). In AT_R_ and PT_R_, the 16 amino acid insertion between β9 and β10 contributes to an ∼25 amino acid ‘lid’, hinging at AT_G550_ and AT_N576_ (PT_P571_ and PT_D596_), that folds back over and occludes the presumed receptor binding surface (Figure 5A, Supplemental Figure 5B). The apex of the loop consists of a helical-turn that lies precisely in the groove where HBEGF is bound in the DT/HBEGF co-crystal structure.

**Figure 5.**
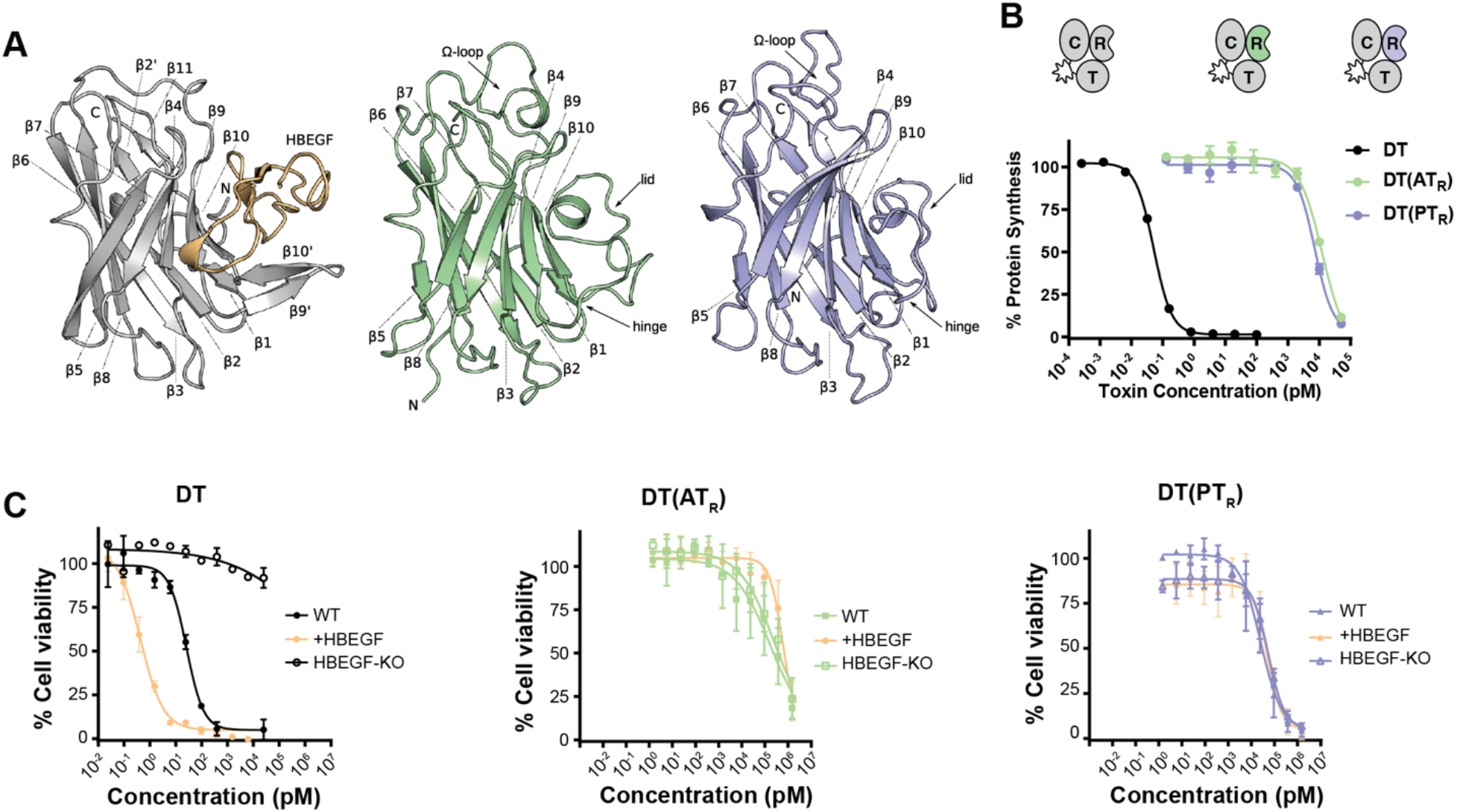
R-domains of AT and PT do not bind HBEGF. (A) R domain structural comparison highlighting conserved β-sandwich motif. The structure of DT_R_ in complex with HBEGF (yellow) was generated from PDB entry 1XDT. The Ω-loop between β8 and β9, and the lid structure between β9 and β10 are labeled. (B) Dose titration of DT and DT R-domain chimeras DT(AT_R_), and DT(PT_R_) on Vero cells (mean ± SD, n=3). (C) Dose titration of DT, DT(AT_R_), and DT(PT_R_) on WT, HBEGF_KO_ and +HBEGF cells (mean ± SD, n=3). DT sensitivity (EC_50_) tracked with expression of HBEGF (WT-26 pM, HBEGF_KO_-N/A, +HBEGF-0.43 pM). Sensitivity to DT(AT_R_) and DT(PT_R_) was unaffected by HBEGF expression (WT-169 nM and 25 nM, HBEGF_KO_-245 nM and 55 nM, +HBEGF-608 nM and 50 nM).

To determine whether AT_R_ and PT_R_ bind and use human HBEGF as a cellular receptor, we generated chimeric constructs with DT in which the R domain of DT was replaced with the R domain from AT or PT. Chimeric DT(AT_R_) and DT(PT_R_) displayed greatly attenuated toxicity toward Vero cells with EC_50_’s of 12 and 7 nM, respectively (Figure 5B). To confirm that the chimeric constructs do not engage HBEGF, we engineered ARN8 cells (that have basal levels of HBEGF) to have either no HBEGF (HBEGF_KO_), or high levels of HBEGF (+HBEGF). Whereas DT sensitivity tracked with expression of HBEGF (Figure 5C), neither DT(AT_R_) nor DT(PT_R_) were affected by knockout or overexpression of HBEGF, further demonstrating that AT and PT do not recognize HBEGF as a cellular receptor. In fact, levels of toxicity for DT(AT_R_) and DT(PT_R_) were indistinguishable for a DT construct devoid of a receptor binding moiety (DT_1-389_) (Supplemental Figure 3B). Attempts to unmask a cryptic HBEGF binding site through genetic deletion of the lid in AT_R_ proved unsuccessful (Supplemental Figure 3C), further demonstrating a lack of receptor recognition and suggesting that the lids are not responsible for the lack of binding to HBEGF. Taken together, these data demonstrate that AT and PT are unable to bind HBEGF or any other human cell-surface protein as a receptor to enter cells, thus playing a major role in the inability of the AT and PT holotoxins to intoxicate human cells.

## Discussion

We have presented a structural and functional analysis of two distant homologs of DT, which has provided novel insights into the molecular evolution of DT and the determinants of host specialization. This study was enabled by the recent discovery of the first distant DT-like relatives (*i*.*e*., all sharing <35% sequence identity with DT) from bacterial lineages outside of *Corynebacterium* with no known association with human hosts. Prior to this, the only known DT homologs were the highly related toxins from *C. ulcerans* and *C. pseudotuberculosis* (95% identity), which were highly toxic to human cells and offered little with respect to understanding the evolutionary history of DT. Moreover, the uniqueness of the structure of DT within the PDB made unravelling the molecular evolution of DT and the adaptations that enabled specialization towards human hosts difficult.

We found that the holotoxins from *S. albireticuli* and *S. peptonophila*, despite being virtually non-toxic to human cells, had strikingly similar structures to DT. Through a comprehensive domain-by-domain analysis, we showed that the factors that contribute to host cell toxicity are demonstrably localized within the B fragments of the toxins (binding and translocation), rather than the A fragments, which displayed comparable cytotoxicity if delivered into the Vero cell cytosol via the DT binding and delivery machinery. That the cytotoxic action of DT_C_ is functionally conserved in these distant homologs likely parallels the highly conserved nature of eEF2 across eukaryotes and is consistent with the observation that DT retains ADP ribosylation activity against yeast eEF-2 [21]. Importantly, these findings strongly suggest that AT and PT are “toxins” in a different context (*i*.*e*., host) where their respective B fragments are optimized to deliver the catalytic fragments into their respective hosts.

Within the catalytic fragment of DT, the residues that have been shown to be important for binding and coordinating NAD^+^ in the active site of DT are well characterized (H21, Y54, Y65, E148) [22-25]. These four residues are conserved within the active sites of other ADP ribosyltransferase enzymes including Exotoxin A, Cholix toxin and PARP. By contrast, in AT and PT, only Y54 and E148 are absolutely conserved: in both, an Arginine is seen in place of the H21 residue; and in AT an Asparagine is seen in place of Y65. Interestingly the pertussis toxin and cholera toxin ADPRT enzymes also have an Arginine at the position equivalent to DT_H21_ (Supplemental Figure 1), indicating at least some functional equivalency in this position between Histidine and Arginine. Future studies on the substrate specificity and enzyme kinetics are needed to understand if these changes confer any additional unique properties to these enzymes.

As might be expected, the receptor binding domains of AT and PT appear to be important determinants of host specificity, with neither able to bind human HBEGF. Without any knowledge of the preferred host of these toxins, we are unable to distinguish at this time whether AT_R_ and PT_R_ bind HBEGF in their respective hosts, an HBEGF-like receptor, or an entirely different receptor. While the ‘lid’ structure is poorly conserved in terms of both sequence and structure (Supplemental Figure 3A), the fact that it is present in both homologs raises several questions. Is the ‘lid’ merely a gate that can move out of the way to accommodate a receptor, or is the surface presented by the ‘lid’ responsible for mediating its receptor binding activity? Also, if this structural feature was present in the common ancestor of DT, AT, and PT, did the loss of this structural feature enable binding to receptors like HBEGF in non β-clade members of the family?

The highly attenuated proteolytic activation of AT and PT between A and B fragments and subsequent impact on toxicity (Figure 3) demonstrates yet another evolutionary lever that can be pulled to tune host specificity. While we cannot exclude cleavage by different proteases in their preferred host organisms, neither appears to be cleavable in Vero cells and this does not affect toxicity to the degree which might be expected. This has similarly been observed with another AB family toxin, TcdB from *Clostridium difficile*, where inactivation of the enzyme releasing domain results in a slight attenuation of toxicity and a temporal delay of the effector function [27]. It is possible that like what has been proposed for TcdB, the catalytic fragments of AT and PT are still translocated into the cytosol, however they remain tethered to the endosome due to the non-cleaving loop joining them to the B fragment. In this scenario, the C domain would have far less freedom of movement within the cell but may nonetheless engage its target (eEF-2). From an evolutionary standpoint, it is unlikely that the formation of the disulfide linkage could occur spontaneously with the furin cleavage motif. The most likely scenario is that the disulfide bond preceded the protease cleavage site, as the inverse order would result in the C domain being released and trapped in the endosome. AT and PT may thus reflect an early diverging lineage of DT-like toxins that lack some of the key characteristics that emerged in the DT lineage.

Understanding how the T-domain of DT evolved the ability to deliver DT_C_ across the endosomal membrane remains an open question for DT along with most other similarly translocating toxins. Uncovering the determinants of translocation has implications for understanding how proteins interact with and cross membranes to cause disease as well as for rational development of intracellular delivery vectors. We found that AT and PT possess marginally functional translocases with a lower pH threshold for unfolding and pore formation in vitro. Sensing of the endosomal pH by titratable residues within the T-domains has long been considered the first step in the process of translocation. Of the three groups of titratable amino acids, only histidine (pKa 6) and the acidic residues, aspartic and glutamic acid (pKa ∼4), are expected to be protonated during endosomal acidification. All other DT-like proteins identified to date contain at least six His residues in their respective T domains [6], presumably serving equivalent roles. The mechanistic role of these six His residues have been studied extensively [17, 19, 20, 26]. The complete lack of His residues in AT_T_ and an altered arrangement of His residues in PT_T_ relative to DT_T_ suggests that initiating the process of unfolding and pore-formation at higher pH’s may lead to more efficient endosomal escape and subsequent translocation in DT (Figure 4D-F). While this more acidic pH transition may truly represent a biologically relevant phenomenon, we cannot rule out the possibility that AT_T_ and PT_T_ are simply less efficient translocases. The lack of His residues in AT_T_ and low number in PT_T_ thus represent a shared feature that may be due to one of two evolutionary scenarios: either the shared precursor of all DT-like proteins contained few or no His residues in its T domain, and His residues were acquired in the α-clade, optimizing the pH transition to escape from the early endosome prior to lysosomal fusion; or DT-like proteins of the β-clade have lost His residues present in the shared ancestral toxin. In other words, these characteristics of AT and PT may be representative of a more ancient lineage of the DT family tree that are simply less efficient translocases, or their preferred hosts occupy a particular niche or possess a particular biology which would make a lower pH transition optimal for endosomal escape.

While other toxins may possess distant homology with regions of DT (*i*.*e*., conserved ADPRT activity), AT and PT represent the first true homologs of DT to be characterized structurally as they are the first to possess a complete DT-like three-domain architecture. This is consistent with our earlier sequence-based analysis, which placed AT and PT among the closest phylogenetic neighbors of DT outside of *Corynebacterium*. Despite widely divergent sequences, both AT and PT display remarkable structural conservation of overall domain architecture as well as within individual domains. We demonstrate that while DT shares several features with distantly related DT-like toxins targeting unknown (non-human) hosts, it also possesses features that are absent from these relatives through which we can potentially trace the emergence of human-specificity, and therefore the origins of the disease diphtheria. More broadly, the field of bacterial toxins has historically assumed a human-centric approach, as toxins have been identified primarily from clinical isolates. With the abundance of easily accessible and ever-expanding genomic databases, a new paradigm for toxin discovery has been enabled.

## Conclusions

Decades of studies on DT and other toxins with intracellular targets that mediate their own entry into cells, such as botulinum neurotoxin, tetanus toxin and anthrax toxin, have helped us to understand in detail how these proteins intoxicate at the molecular and cellular level to ultimately cause disease. By contrast, the origins of these toxins and their adaptation to the human host are fundamental questions that have not been readily addressable owing to the paucity of distant homologs adapted to different hosts to compare to. In this work, through the structural and functional characterization of two recently identified distant homologs of DT not adapted to human hosts, we uncovered insights into the evolution of DT and determinants of specificity to humans. We found that unlike other bacterial toxin families that have slight variations in enzyme function and intracellular targets, the DT-like family of proteins all conserved their catalytic domain function and target. We also found that much like how viruses jump from one host species to another through a series of stepwise evolutionary events that enable virus entry and replication, DT, and perhaps bacterial toxin specificity, for humans appears to derive from adaptations to host factors implicated in the entry process (i.e., receptor binding and subsequent internalization and enzyme release). Whereas receptor-switching through alterations in the receptor-binding surface may be sufficient to change tissue tropism within a given species (as is the case with the Large Clostridial Toxin family [32]), our work here seems to suggest that adaptation to multiple host-specific factors and processes implicated in toxin uptake (*i*.*e*., receptors, endosomal pH, activating proteases, etc.) are required to produce maximal toxicity toward a given host. Finally, this work demonstrates the importance of structural plasticity in the evolution of not just DT and DT-family proteins, but perhaps all bacterial toxins.

## Materials and Methods

### Chimera Generation and Protein Purification

*E. coli* codon optimized gBlocks® gene fragments for full-length DT, AT and PT were purchased from Integrated DNA Technologies (IDT). Primary amino sequences were obtained for AT and PT from NCBI database entries WP_095582082.1 and WP_073156187.1, respectively. Individual domains were amplified by PCR, and domain-swapped chimeras were generated with the NEBuilder® HiFi DNA Assembly Cloning Kit (New England BioLabs). All toxins and chimeras were expressed and purified with cleavable N-terminal 6His-SUMO fusions (ThermoFisher), grown to an OD_600_ of 0.8 in LB media, and induced for 18 hours at 18°C using 0.1 mM IPTG. Cell pellets were pelleted by centrifugation and then re-suspended in lysis buffer (20 mM Tris pH 8.0, 500 mM NaCl, 1% protease inhibitor cocktail (Sigma), 1 mg/mL lysozyme, 0.01% benzonase (Sigma)) and lysed by 3-passes through an Emulsiflex C3 (Avestin) at 15,000 psi. Cell lysates were clarified by centrifugation (20 minutes at 18,000x g) and bound to a 5 mL HisTrap™ FF Crude column (GE Healthcare) and eluted with 50-75 mM imidazole. Eluted protein was diluted to ∼150 mM NaCl and incubated overnight at 4°C with SUMO protease to cleave the affinity tag. Cleaved protein was separated from SUMO, SUMO protease and uncleaved protein with Ni-NTA resin, concentrated by centrifugation and exchanged into 20 mM Tris pH 7.5, 150 mM NaCl by dialysis.

### Protein Synthesis Inhibition

Vero cells (ATCC) were transduced with a lentivirus expressing NanoLuc® luciferase (Vero-NLucP cells) as described previously [8]. Vero-NLucP cells were plated (5,000 cells/well) in a white clear-bottom 96-well plate and treated with toxin the following day. Following overnight incubation (20 hours), luminescence signal was developed using the NanoGlo® Luciferase Assay kit (Promega) and read with a Spectramax M5e plate reader (Molecular Devices). Relative luminescence units (RLU) were normalized relative to untreated cells and data was analyzed with GraphPad Prism 7.0. For protein synthesis inhibition assays in HEK293T cells, cells were transiently transfected with the pNL3.2CMV (Promega) rather than by lentiviral transduction. Cells were plated in a 6-well plate at 1×10^6^ cells/well and transfected the following day with 3.0 µg of plasmid DNA, in a 3:1 ratio of FuGENE^®^HD (Promega) transfection reagent to DNA. After 24 hours, cells were re-plated at 5×10^3^ cells/well, in a white clear-bottom 96-well plate and treated the following day with toxin. After overnight incubation (20 hours), luminescence signal was developed using the NanoGlo® Luciferase Assay kit (Promega) similar to above. Wildtype and DPH4^-/-^ HEK293T cells were a gift from Dr. Mikko Taipale at the University of Toronto.

### Cell Viability

Cells (Vero-NLucP, HEK293T, ARN8) were plated in a black clear-bottom 96-well plate (5×10^3^ cells/well) and treated with toxin the following day. After 48 hours, cells were treated with PrestoBlue™ reagent (Thermo Fisher Scientific), incubated for 2 hours, and fluorescence was read (555 nm excitation and 585 nm emission) with a Spectramax M5 plate reader (Molecular Devices). Fluorescence measurements were normalized relative to untreated cells and data was analyzed with GraphPad Prism 7.0.

### Crystallization

All crystals were grown by hanging-drop vapor diffusion at 20°C. For AT, 1 µL of selenomethionine derivatized AT (11.8 mg/mL) was mixed with 1 µL of mother liquor (10 mM Tris-HCl pH 7.0, 200 mM calcium acetate hydrate, 20% PEG3000) and dehydrated over 200 µL of mother liquor. Diffraction quality AT crystals were obtained following successive rounds of micro-seeding and flash frozen in liquid nitrogen without any additional cryoprotectant. PT crystals were grown from drops containing 1 µL PT (15.8 mg/mL) and 1 µL mother liquor (100 mM potassium iodide, 22% PEG3350) dehydrated over 200 µL of mother liquor. PT crystals used for data collection were grown following successive rounds of micro-seeding in drops dehydrated over 550 mM ammonium sulfate, and flash frozen in liquid nitrogen without any additional cryoprotectant. Datasets were collected remotely at a wavelength of 0.979 Å for AT crystals and 2.0 Å for PT crystals on the AMX beamline at NSLSII (Brookhaven National Labs).

### Structure Solution

All diffraction data was processed with XDS using the AutoPROC package [28]. Structure solution of AT by SAD phasing was carried using the CRANK2 pipeline [29] in CCP4 [30]. The initial solution yielded 39 heavy atom sites (selenium) with an FOM of 56.8, producing a partial model with an R_factor_ of 43.37% (Buccaneer). A single monomer from the partial model was used to perform molecular replacement using the PHENIX software package[31], which located 4 copies of AT in the asymmetric unit, which were then re-built with PHENIX-AutoBuild. SAD phasing of PT was performed with PHENIX-AutoSol, yielding 18 heavy atom sites (iodine) with an FOM of 35.2 and a partial model (monomer) with an R_factor_ of 48.6%. Model building and refinement was carried out through multiple iterations of manual building in Coot[32] and PHENIX-Refine until R and R_free_ values converged and geometry statistics reached suitable ranges.

### Sequence alignments

Sequence identity and similarity between DT, AT, and PT were calculated from a multiple sequence alignment generated with the T-coffee server using the M-coffee algorithm (http://tcoffee.crg.cat/) [33].

### Phylogenetic Tree Construction

In order to compare DTs and their homologs phylogenetically, a set of DTs from *Corynebacterium* spp. were combined DT homolog proteins from various species of Actinobacteria. The set of sequences used here is the same as previously identified by Mansfield et al. 2018 [6], with the addition of 5 additional DT homolog sequences (WP_116115734.1 from *Austwickia chelonae*, WP_169314017.1 from *Streptomyces pinterrae*, WP_136737859.1 from *Streptomyces* sp. jys28, WP_120757473.1 from *S. klenkii*, and WP_189053160.1 from *Longimycelium tulufanense*). Multiple sequence alignments of full-length proteins were created using MAFFT (version 7.487; the high-accuracy L-INS-i algorithm) [34], ClustalO (version 1.2.4) [35], MUSCLE (version 3.8.1551) [36], and PRANK (version v.170427; with the -F flag to maintain insertions over alignment iterations) [37]. Maximum likelihood phylogenetic trees were generated for each of these alignments with RAxML (version 8.2.12) [38]. Each RAxML tree used rapid bootstrapping followed by a slow maximum likelihood search, the autoMRE bootstopping algorithm, automatic amino acid substitution matrix selection, and the GAMMA model of rate heterogeneity. The tree that yielded the highest likelihood was selected as the representative topology. The topology of the RAxML tree was further supported through Bayesian phylogenetic inference with MrBayes (version 3.2.7)[39]. MrBayes was run with the GTR model and gamma-distributed rate heterogeneity, the “mixed” amino acid substitution matrix (allowing the program to sample from the available matrices and select the best one), 3 runs with random starting trees and a chain length of 1,000,000 generations, automatic simulation stopping via average standard deviation of split frequencies, and a 25% burn-in fraction. The posterior support of clades of the resulting consensus tree are depicted in Figure 1. All scripts needed to reproduce the bioinformatic analysis can be found at https://github.com/mjmansfi/Sugiman-Marangos_Gill_etal_2021.

### Vero Lysate Processing Assay

Vero-NLucP cells (3×10^6^) were resuspended in 500 µl lysis buffer (0.05% Triton X100 in PBS). 2.5 µg of toxin was incubated with 0.5 µL of cell lysate overnight at 37°C in a total volume of 5 µL PBS with 2 mM CaCl_2_. The reaction was stopped by adding 5 µL of 2X SDS-PAGE loading dye. Samples were boiled for 3 minutes and separated on a 4-20% precast polyacrylamide gel (Biorad).

### TNS Fluorescence Assay

2-(p-toluidino) naphthalene6-sulphonic acid (TNS) assays were performed as described previously [40]. Briefly, proteins (20 nM final concentration) were incubated in 150 mM citrate phosphate buffers ranging from pH 3.0 to 7.5 with 150 µM 2-(*p*-Toluidinyl) naphthalene-6-sulfonic acid, sodium salt (Invitrogen). Following 25-minute incubation at 37°C, fluorescence was read (366 nm excitation and 400-500 nm emission scan) with a Spectramax M5 plate reader (Molecular Devices).

### DSF Assay

Differential scanning fluorometry (DSF) was performed as described previously[41]. Proteins were diluted to a final concentration of 0.1 mg/mL in 150 mM citrate phosphate buffers ranging from pH 3.0 to 7.5 containing 5x SYPRO Orange (Invitrogen). SYPRO Orange fluorescence was monitored over a 25°C to 80°C temperature gradient (30 s increments) using a BioRad CFX96 qRT-PCR thermocycler.BioRad CFX Manager 3.1 was used to integrate fluorescence curves and calculate melting temperatures.

### Liposomal dye release assay

Unilamellar liposomes (DOPC, 0.8% DGS-NTA, Avanti Polar Lipids) were prepared as previously described [5]. Briefly, 1,2-dioleoyl-sn-glycero-3-phosphocholine (DOPC) (Avanti Polar Lipids) was combined with 0.8% 1,2-dioleoyl-sn-glycero-3-[(N-(5-amino-1-carboxypentyl)iminodiacetic acid)succinyl] (nickel salt) (DGS-NTA[Ni]) (Avanti Polar Lipids), dried with N_2_ and incubated for 1 hour in a vacuum desiccator. Lipids were resuspended in 20mM Tris pH 8, 35mM 8-Hydroxypyrene-1,3,6-trisulfonic acid (HPTS), 50mM p-xylene-bis-pyridinium bromide (DPX) (Thermo Fischer) and subject to 10X freeze-thaw cycles in dry ice and 42°C water bath, and 10X extrusions using a 200 µm filter. The liposomes were then purified by gel filtration using a Hi Prep 16/60 Sephacryl S-300 HR column (GE Healthcare) in 150 mM NaCl, 20 mM Tris pH 8 buffer. Proteins were added in a ratio of 1:10,000 with liposomes, with a final liposome concentration of ∼400 µM, in 150 mM citrate phosphate buffer ranging from pH 4.0 to 7.5, in 0.5 pH intervals. Assays were done in 96-well opaque plates (Corning), and fluorescence was monitored over a 20-minute interval, with readings being taken every 30 seconds (excitation 403nm, emission 510nm). Data were normalized to % of total HPTS fluorescence, by adding 0.3% Triton X-100.

### Generation of bulk ARN8 HBEGF_KO_ cell line

crRNA targeting the HBEGF (5’ - ACGGACCAGCTGCTACCCCT) was designed using the Integrated DNA Technologies (www.idtdna.com) design tool. The gRNA:Cas9 ribonucleoprotein complex was assembled according to the manufacturer’s protocol (Integrated DNA Technologies) and reverse transfected using Lipofectamine RNAiMAX (Thermo Fisher Scientific) into ARN8 cells (3×10^5^ cells in a 6-well plate). Following 48 hours of incubation, 10 nM of WT DT was added to the plates to select for HBEGF knockout cells.

### hHBEGF overexpression in ARN8 HBEGF_KO_ cells

The gene for human HBEGF was synthesized and cloned into the MCS of PiggyBac vector PB-CMV-MCS-EF1α-Puro (System Biosciences) by restriction digest with EcoRI and BamHI (pCMV-hHBEGF). ARN8 HBEGF_KO_ cells were transfected at 80% confluence in a 6-well plate with pCMV-hHBEGF (2 µg) and PB Transposase vector (0.5 µg) (System Biosciences) using FuGENE® HD (Promega) as per the manufacturer’s protocol. Selection was performed with puromycin (6 µg/mL) for 1 week beginning 48 hours after transfection.

## Acknowledgments

This project was supported in part by a National Sciences and Engineering Research Council of Canada Discovery Grant (RGPIN 2017 06817) and the Canadian Institutes of Health Research Operating Grant (R.M.) The authors wish to thank Dr. R. John Collier for his expert guidance throughout this project and for critically reading and providing feedback on this manuscript. HEK293T and HEK293T DPH4_KO_ cells were obtained as a generous gift from Dr. Mikko Taipale (University of Toronto). M.J.M. gratefully acknowledges funding from the Japan Society for the Promotion of Science as a JSPS International Research Fellow (Luscombe Unit, Okinawa Institute of Science and Technology Graduate University).

## References

1. Collier, R.J., Understanding the mode of action of diphtheria toxin: a perspective on progress during the 20th century. Toxicon, 2001. 39(11): p. 1793–803.

2. Doxey, A.C., M.J. Mansfield, and C. Montecucco, Discovery of novel bacterial toxins by genomics and computational biology. Toxicon, 2018. 147: p. 2–12.

3. Mansfield, M.J., J.B. Adams, and A.C. Doxey, Botulinum neurotoxin homologs in non-Clostridium species. FEBS Lett, 2015. 589(3): p. 342–8.

4. Visschedyk, D., et al., Certhrax toxin, an anthrax-related ADP-ribosyltransferase from Bacillus cereus. J Biol Chem, 2012. 287(49): p. 41089–102.

5. Orrell, K.E., et al., The C. difficile toxin B membrane translocation machinery is an evolutionarily conserved protein delivery apparatus. Nat Commun, 2020. 11(1): p. 432.

6. Mansfield, M.J., et al., Identification of a diphtheria toxin-like gene family beyond the Corynebacterium genus. FEBS Lett, 2018. 592(16): p. 2693–2705.

7. Holm, L., DALI and the persistence of protein shape. Protein Sci, 2020. 29(1): p. 128–140.

8. Park, M., et al., Intracellular Delivery of Human Purine Nucleoside Phosphorylase by Engineered Diphtheria Toxin Rescues Function in Target Cells. Mol Pharm, 2018.

9. Middlebrook, J.L., R.B. Dorland, and S.H. Leppla, Association of diphtheria toxin with Vero cells. Demonstration of a receptor. J Biol Chem, 1978. 253(20): p. 7325–30.

10. Tsuneoka, M., et al., Evidence for involvement of furin in cleavage and activation of diphtheria toxin. J Biol Chem, 1993. 268(35): p. 26461–5.

11. Williams, D.P., et al., Cellular processing of the interleukin-2 fusion toxin DAB486-IL-2 and efficient delivery of diphtheria fragment A to the cytosol of target cells requires Arg194. J Biol Chem, 1990. 265(33): p. 20673–7.

12. Choe, S., et al., The crystal structure of diphtheria toxin. Nature, 1992. 357(6375): p. 216–22.

13. Rosconi, M.P., G. Zhao, and E. London, Analyzing topography of membrane-inserted diphtheria toxin T domain using BODIPY-streptavidin: at low pH, helices 8 and 9 form a transmembrane hairpin but helices 5-7 form stable nonclassical inserted segments on the cis side of the bilayer. Biochemistry, 2004. 43(28): p. 9127–39.

14. Wang, J. and E. London, The membrane topography of the diphtheria toxin T domain linked to the a chain reveals a transient transmembrane hairpin and potential translocation mechanisms. Biochemistry, 2009. 48(43): p. 10446–56.

15. Johnson, V.G., et al., The role of proline 345 in diphtheria toxin translocation. J Biol Chem, 1993. 268(5): p. 3514–9.

16. Zhan, H., et al., Effects of mutations in proline 345 on insertion of diphtheria toxin into model membranes. J Membr Biol, 1999. 167(2): p. 173–81.

17. Ladokhin, A.S., pH-triggered conformational switching along the membrane insertion pathway of the diphtheria toxin T-domain. Toxins (Basel), 2013. 5(8): p. 1362–80.

18. Leka, O., et al., Diphtheria toxin conformational switching at acidic pH. FEBS J, 2014. 281(9): p. 2115–22.

19. Rodnin, M.V., et al., Conformational switching of the diphtheria toxin T domain. J Mol Biol, 2010. 402(1): p. 1–7.

20. Rodnin, M.V., et al., The pH-Dependent Trigger in Diphtheria Toxin T Domain Comes with a Safety Latch. Biophys J, 2016. 111(9): p. 1946–1953.

21. Mateyak, M.K. and T.G. Kinzy, ADP-ribosylation of translation elongation factor 2 by diphtheria toxin in yeast inhibits translation and cell separation. J Biol Chem, 2013. 288(34): p. 24647–55.

22. Papini, E., et al., Histidine 21 is at the NAD+ binding site of diphtheria toxin. J Biol Chem, 1989. 264(21): p. 12385–8.

23. Papini, E., et al., Histidine-21 is involved in diphtheria toxin NAD+ binding. Toxicon, 1990. 28(6): p. 631–5.

24. Tweten, R.K., J.T. Barbieri, and R.J. Collier, Diphtheria toxin. Effect of substituting aspartic acid for glutamic acid 148 on ADP-ribosyltransferase activity. J Biol Chem, 1985. 260(19): p. 10392–4.

25. Blanke, S.R., K. Huang, and R.J. Collier, Active-site mutations of diphtheria toxin: role of tyrosine-65 in NAD binding and ADP-ribosylation. Biochemistry, 1994. 33(51): p. 15494–500.

26. Perier, A., et al., Concerted protonation of key histidines triggers membrane interaction of the diphtheria toxin T domain. J Biol Chem, 2007. 282(33): p. 24239–45.

27. Li, S., et al., Cytotoxicity of Clostridium difficile toxin B does not require cysteine protease-mediated autocleavage and release of the glucosyltransferase domain into the host cell cytosol. Pathog Dis, 2013. 67(1): p. 11–8.

28. Vonrhein, C., et al., Data processing and analysis with the autoPROC toolbox. Acta Crystallogr D Biol Crystallogr, 2011. 67(Pt 4): p. 293–302.

29. Skubak, P. and N.S. Pannu, Automatic protein structure solution from weak X-ray data. Nat Commun, 2013. 4: p. 2777.

30. Winn, M.D., et al., Overview of the CCP4 suite and current developments. Acta Crystallogr D Biol Crystallogr, 2011. 67(Pt 4): p. 235–42.

31. Adams, P.D., et al., PHENIX: a comprehensive Python-based system for macromolecular structure solution. Acta Crystallogr D Biol Crystallogr, 2010. 66(Pt 2): p. 213–21.

32. Emsley, P. and K. Cowtan, Coot: model-building tools for molecular graphics. Acta Crystallogr D Biol Crystallogr, 2004. 60(Pt 12 Pt 1): p. 2126–32.

33. Notredame, C., D.G. Higgins, and J. Heringa, T-Coffee: A novel method for fast and accurate multiple sequence alignment. J Mol Biol, 2000. 302(1): p. 205–17.

34. Katoh, K. and D.M. Standley, MAFFT multiple sequence alignment software version 7: improvements in performance and usability. Mol Biol Evol, 2013. 30(4): p. 772–80.

35. Sievers, F., et al., Fast, scalable generation of high-quality protein multiple sequence alignments using Clustal Omega. Mol Syst Biol, 2011. 7: p. 539.

36. Edgar, R.C., MUSCLE: multiple sequence alignment with high accuracy and high throughput. Nucleic Acids Res, 2004. 32(5): p. 1792–7.

37. Loytynoja, A. and N. Goldman, An algorithm for progressive multiple alignment of sequences with insertions. Proc Natl Acad Sci U S A, 2005. 102(30): p. 10557–62.

38. Stamatakis, A., RAxML version 8: a tool for phylogenetic analysis and post-analysis of large phylogenies. Bioinformatics, 2014. 30(9): p. 1312–3.

39. Ronquist, F., et al., MrBayes 3.2: efficient Bayesian phylogenetic inference and model choice across a large model space. Syst Biol, 2012. 61(3): p. 539–42.

40. Lanis, J.M., S. Barua, and J.D. Ballard, Variations in TcdB activity and the hypervirulence of emerging strains of Clostridium difficile. PLoS Pathog, 2010. 6(8): p. e1001061.

41. Tam, J., et al., Host-targeted niclosamide inhibits C. difficile virulence and prevents disease in mice without disrupting the gut microbiota. Nat Commun, 2018. 9(1): p. 5233.

