## Supplemental Material for "Structure-function analysis of distant diphtheria toxin homologs reveals insights into host adaptation"

**Supplemental Table 1. Data collection and refinement statistics.**

| | **Albireti Toxin (AT) PDB ID 7RI3** | **Peptono Toxin (PT) PDB ID 7RB4** |
|---|---|---|
| | **Data Collection** | |
| **Wavelength (Å)** | 0.9795 | 2.00 |
| **Resolution (Å)** | 60.8-2.69 (2.696-2.687) | 48.14-2.19 (2.192-2.185) |
| **Space group** | $P2_12_12$ | $P2_12_12_1$ |
| **Unit cell – a, b, c (Å)** | 106.95, 137.08, | 77.65, 82.46, 91.82 |
| **α, β, γ (°)** | 203.51 90, 90, 90 | 90, 90, 90 |
| **Total reflections** | 1,168,104 (11,867) | 333,179 (560) |
| **Unique reflections** | 83,962 (803) | 28,929 (168) |
| **Multiplicity** | 13.9 (14.8) | 11.5 (3.3) |
| **Completeness (%)** | 99.9 (100.0) | 92.7 (54.0) |
| **Mean I/sigma(I)** | 10.9 (2.1) | 16.4 (1.9) |
| **R-merge** | 18.9 (143.5) | 11.8 (58.7) |
| **R-pim** | 5.2 (38.3) | 3.5 (35.3) |
| **$CC_{1/2}$** | 99.8 (83.3) | 99.6 (66.3) |
| **Wilson B-factor** | 49.43 | 28.59 |
| | **Structure Refinement** | |
| **Reflections** | 83,614 (8,257) | 28,814 (1,832) |
| **R-work/R-free** | 20.24 / 24.75 | 20.15 / 22.50 |
| **Number of atoms** | 18,286 | 5,080 |
| **macromolecules** | 17,703 | 4,662 |
| **ligands** | 156 | 26 |
| **solvent** | 511 | 392 |
| **RMS bonds (Å)** | 0.006 | 0.002 |
| **RMS angles (°)** | 0.79 | 0.50 |
| **Ramachandran allowed (%)** | 99.87 | 100.00 |
| **Average B-factor** | 52.13 | 33.38 |
| **macromolecules** | 52.48 | 33.08 |
| **ligands** | 56.56 | 61.03 |
| **solvent** | 39.61 | 35.15 |

Statistics for the highest-resolution shell are shown in parentheses.

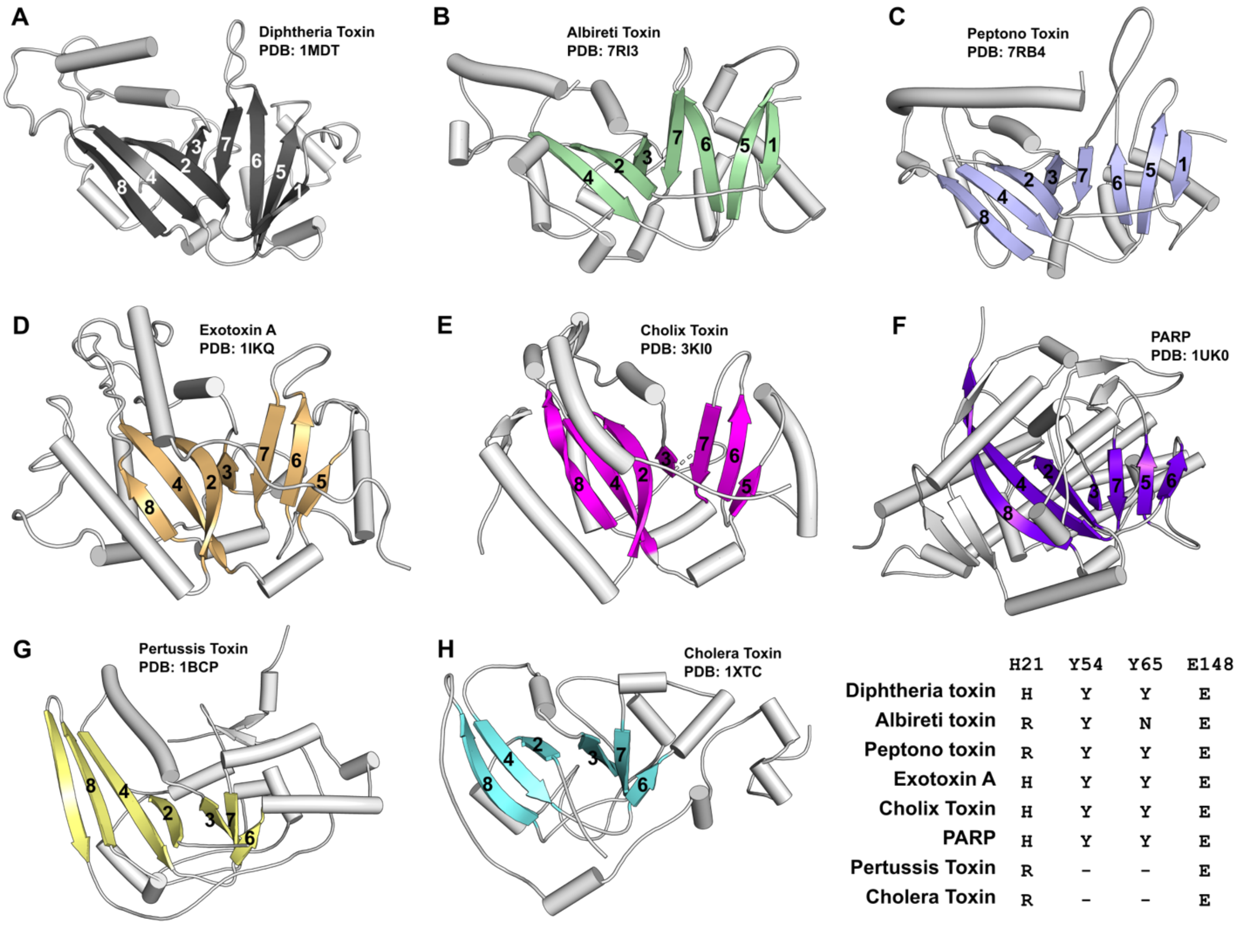


| | H21 | Y54 | Y65 | E148 |
|---|---|---|---|---|
| Diphtheria toxin | H | Y | Y | E |
| Albireti toxin | R | Y | N | E |
| Peptono toxin | R | Y | Y | E |
| Exotoxin A | H | Y | Y | E |
| Cholix Toxin | H | Y | Y | E |
| PARP | H | Y | Y | E |
| Pertussis Toxin | R | - | - | E |
| Cholera Toxin | R | - | - | E |

**Supplemental Figure 1. ADP ribosylating enzyme folds.** Conservation of the split β-sheet amongst ART family catalytic domains.

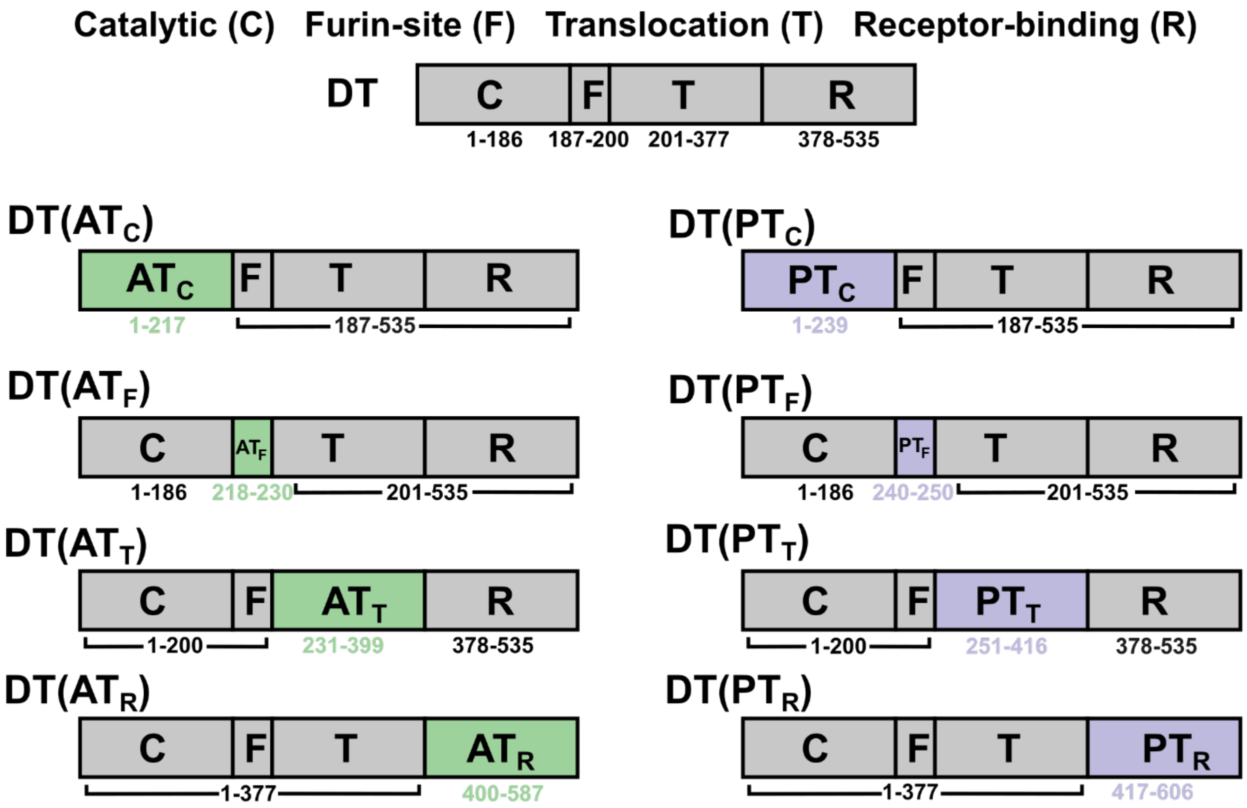


**Supplemental Figure 2. DT chimeras.** Architecture and residue numbering of DT domain swapped chimeras.

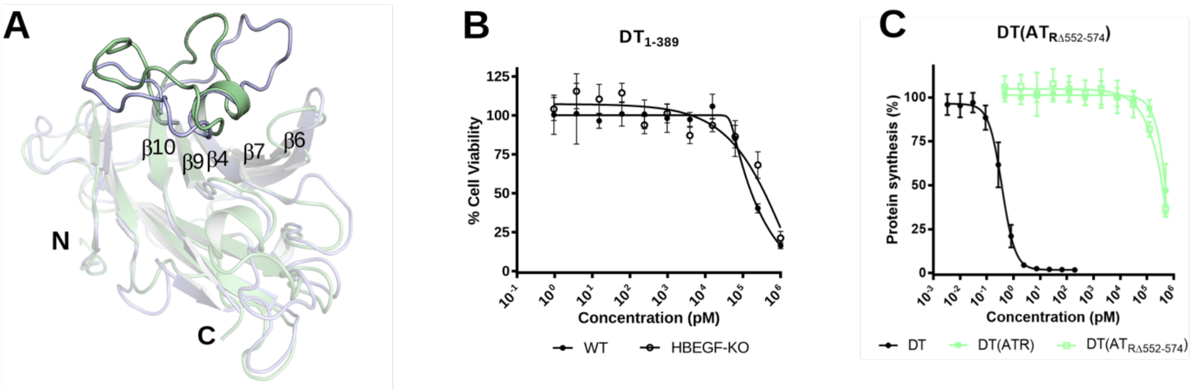


**Supplemental Figure 3. R domain.** (A) Structural differences in the lid structure present in AT and PT. In $PT_R$ the 'lid' is more splayed, and the apex of the loop does not possess as much helical character while in $AT_R$, it adopts a full helical-turn. Sequence conservation between the two proteins in this region is low (11% identical, 22% similar) relative to the full domain (28% identical, 42% similar). (B) Dose titration of $DT_{1-389}$ on WT and $HBEGF_{KO}$ cells (mean ± SD, n=3). Sensitivity to $DT_{1-389}$ was unaffected by HBEGF expression ($EC_{50}$'s: WT-207 nM, $HBEGF_{KO}$-498 nM). (C) The 'lid' sequence in $AT_R$ (residues 552-574) was replaced with two Gly residues in the $DT(AT_R)$ chimeric construct. Dose titration of DT, $DT(AT_R)$, and $DT(AT_{R\Delta 552-574})$ on Vero cells (mean ± SD, n=3).
